# The role of electrostatic interactions in the phase separation of HP1α and its protein binding partners

**DOI:** 10.64898/2026.07.06.736852

**Authors:** Cheenou Her, Rushabh Bhakta, Tonkhla Dankul, Tien M. Phan, Lannah S. Abasi, Jeetain Mittal, Galia T. Debelouchina

## Abstract

Heterochromatin protein 1α (HP1α) is an intrinsic component of heterochromatin domains where it is involved in a diverse set of functions including heterochromatin spreading and organization, chromatin compaction and transcriptional silencing. It has been suggested that HP1α functions through a phase separation mechanism, a process that has been observed *in vitro* in the presence of N-terminal phosphorylation, nucleic acids and nucleosome arrays. HP1α can also interact with numerous binding partners that contain a specific motif called an HP1 access code (HAC). HACs recognize and bind to an interface formed by the chromoshadow (CSD) domains in the HP1α homodimer, the functional form of the protein. It has been shown that some HP1α binding partners can enhance its phase separation ability while others disrupt the process. Here, we focus on the interactions between HP1α and three binding partners, namely the p150 subunit of the chromatin assembly factor 1 (CAF-1), the N-terminal domain of the lamin B receptor (LBR), and the mitotic protein Shugoshin 1 (Sgo1). Using phase separation assays, we show that CAF-1 prevents HP1α phase separation while LBR and Sgo1 enhance it. Binding assays, mutational studies, NMR spectroscopy and computational analysis allow us to dissect the contributions of the HAC motifs, the charge patterns of the binding partner sequences and the role of N-terminal phosphorylation on HP1α in condensate formation. Our results demonstrate that each binding partner uniquely balances these contributions to modulate the properties of HP1α, while electrostatic interactions dominate the regulation of phosphorylated HP1α. These results suggest that HP1α’s binding partners play an important role in the modulation of its properties and the regulation of its functions in distinct biological contexts.

**Statement of significance:** Cellular nuclei are organized into euchromatin and heterochromatin domains, encompassing actively transcribed genes or transcriptionally silent regions, respectively. The regulation of these domains is an active area of study with vast consequences for human health and disease. Here, we focus on the HP1α protein, a key component of heterochromatin compartments that is believed to function through a phase separation mechanism. HP1α interacts with a multitude of binding partners that recognize a specific interaction surface on the protein. While these specific interactions are important, our results suggest that the overall electrostatic network that forms between the proteins has a profound effect on the properties of HP1α. These results have important implications for understanding how HP1α function is regulated in heterochromatin environments.

## Introduction

Heterochromatin protein 1α (HP1α) is a key component of transcriptionally silent constitutive heterochromatin domains that form at telomeres, centromeres and the nuclear periphery.^1^ In a process known as heterochromatin spreading, HP1α recognizes and binds to histone H3 Lys9 di-or trimethylation (H3K9me2/3) on nucleosomes and further interacts with methyltransferases such as SUV39H1/2 that are responsible for the installation of this heterochromatin mark.^2^ This creates a positive feedback loop that allows for the establishment, expansion and maintenance of heterochromatin domains.^1–3^ Furthermore, HP1α can interact with nucleosomes through H3K9me2/3 and through contacts with DNA, a process that promotes chromatin compaction.^4–9^ More recently, HP1α has been shown to form phase separated condensates *in vitro* and, while actively debated in the literature, condensate formation has been suggested to play a role in the formation and function of heterochromatin domains.^10–16^ In humans, there are two additional paralogs, HP1β and HP1γ, which have some overlapping functions with HP1α but differ in biophysical properties (e.g. condensate formation) and distribution throughout the nucleus.^7,13,17,18^ While disease-relevant mutations in the HP1 proteins are rare, changes in expression levels and mis-localization can cause genomic instability and have been linked to cancer and neurodegeneration.^19,20^

HP1α consists of two structured domains, the chromodomain (CD) that contains a binding pocket for H3K9me2/3, and the chromoshadow domain (CSD) that is responsible for homodimerization (**Fig. 1a**).^17^ In addition, there are three disordered regions called the N-terminus extension (NTE), the hinge region, and the C-terminus extension (CTE). The NTE is constitutively phosphorylated in cells, while *in vitro* this modification significantly enhances the phase separation properties of the protein.^10,21–24^ The hinge region, on the other hand, is positively charged and is responsible for interactions with nucleic acids.^7,16,25,26^ It is believed that HP1α functions as a homodimer in cells and that the multivalency promoted by dimerization is required for condensate formation and chromatin compaction.^10,27^ Furthermore, dimerization leads to the creation of a new interacting surface at the base of the CTE regions of the two monomers (**Fig. 1b**),^28^ and this interaction surface acts as a binding platform for many proteins, including SUV39H1/2,^29^ lamin B receptor (LBR),^30^ chromatin assembly factor 1 (CAF-1)^31^ and Shugoshin 1 (Sgo1)^32^. Recognition of a diverse set of binding partners (BPs) is achieved through the so-called PXVXL or PXVXL-like motif on the partner where X can be any amino acid.^33^ Recent bioinformatics analysis, however, has expanded the definition of the recognition motif to include additional residues and to relax the constraints on the P and L positions.^34^ These updated binding motifs have been termed HACs (HP1 access codes)^34^, which is the definition that we adopt here.

**Figure 1.**
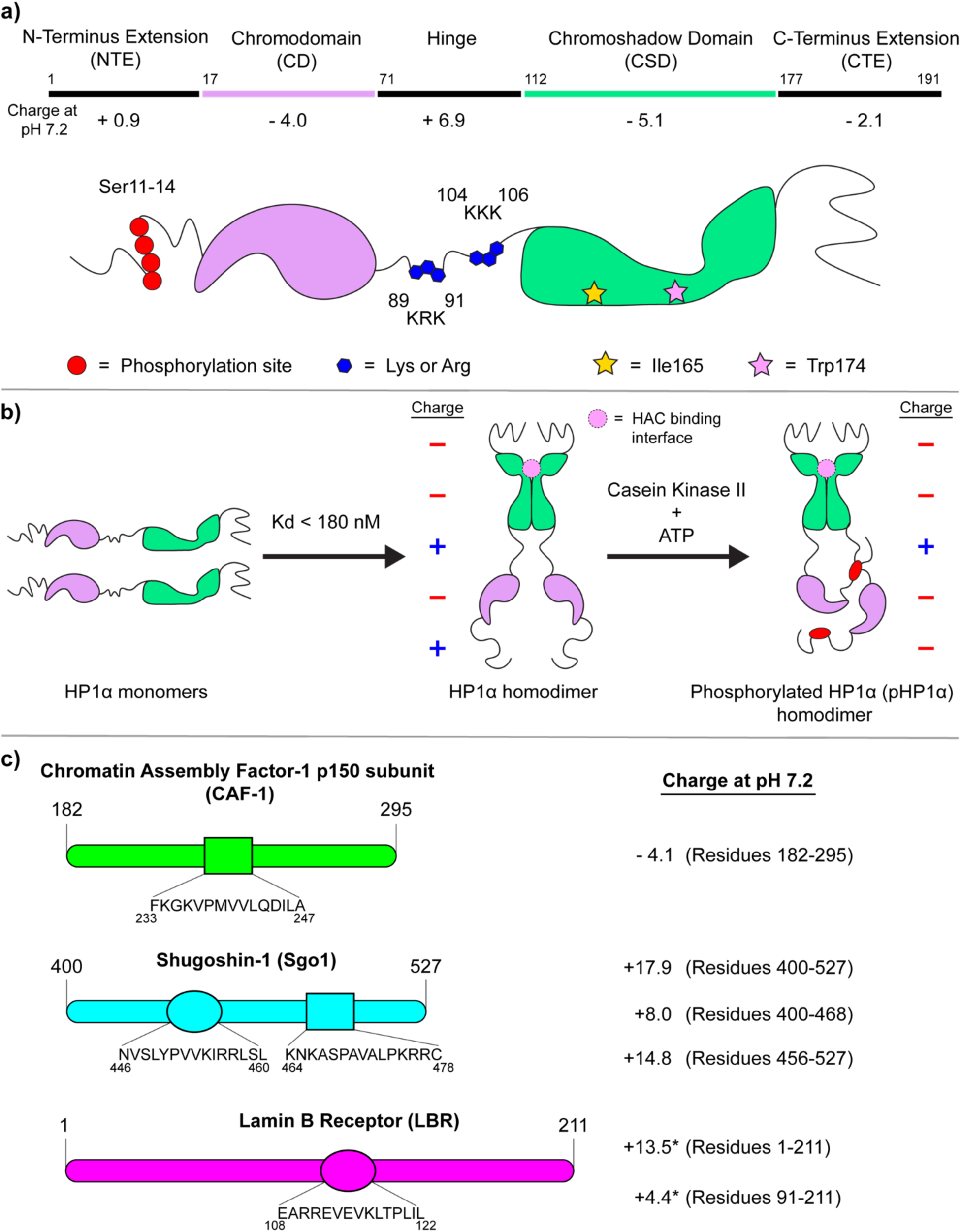
Properties of HP1α and BPs. **a)** Domain organization of HP1α. The net charge of each region was calculated at pH 7.2. The red circles denote serine residues 11-14 that can be phosphorylated; the blue pentagons denote lysine and arginine clusters in the hinge; the gold star indicates Ile165 which is important for dimerization; the pink star denotes Trp174 which is important for BP interactions. **b)** The dimerization of two HP1α monomers leads to the formation of a new interaction surface where BPs can interact through their HAC motifs. The dimer can be further phosphorylated on the NTE by casein kinase II. **c)** BP constructs used in this study. The HAC motifs and the net charge at pH 7.2 are indicated for each construct. Note that the calculated charge for LBR includes the hexahistidine tag sequence (denoted by *).

The fact that HP1α can interact with a diverse set of BPs through a relatively promiscuous binding motif raises the question of how these proteins influence the biophysical properties and functions of HP1α. Previous literature suggests that HP1α co-localizes with many of its BPs in cells and co-partitions with some of them *in vitro* (e.g. SUV39H1/2, TRIM28 and MeCP2).^35,36^ Studies with short BP segments, on the other hand, suggest that some BP peptides can dramatically enhance the ability of HP1α or phosphorylated HP1α (pHP1α) to form condensates while others have a disruptive effect.^10,23^ For example, a positively charged peptide segment of Sgo1 that comprises the relevant HAC significantly lowers the saturation concentration of pHP1α required for phase separation *in vitro*^10,23^, while a negatively charged HAC-containing peptide derived from CAF-1 prevents condensation^23^. The K_d_ values describing the HAC interactions for these peptides are very similar (0.1 – 0.4 μM), suggesting that charge may play a prominent role in the condensation mechanism.^23^

Despite these observations, there is currently no unifying model of how BPs regulate HP1α condensation and how this process balances specific interactions through the HAC-CSD binding interface and electrostatic interactions that promote more transient contacts between other regions of the proteins. Building on our previous work with small BP peptides and pHP1α,^23^ here we undertake a systematic investigation of BP properties that influence the phase separation behavior of HP1α in the presence and absence of N-terminal phosphorylation. In particular, we focus on the following proteins (**Fig. 1c**, **Fig. S1**): 1) a segment comprising residues 182 – 295 in the p150 subunit of CAF-1, a protein involved in nucleosome assembly and DNA damage repair^37–40^; 2) a segment representing residues 1 – 211 of the LBR protein, which is responsible for tethering heterochromatin domains to the nuclear lamina;^41^ and 3) three segments encompassing residues 400 – 527 of Sgo1, a centromeric protein that is crucial for proper chromosome segregation during cell division.^42^ These BP segments span an overall charge from – 4.1 to + 17.9 and represent the diverse HP1α BP functional landscape. We use binding assays, mutational studies, fluorescence imaging, NMR spectroscopy, and molecular simulations to dissect the contributions of HAC-CSD interactions, electrostatic contacts and phosphorylation in HP1α condensate formation. Our results reveal that each BP uniquely balances these contributions to modulate the properties of HP1α, further expanding the interaction and functional landscape of this small but versatile protein.

## Materials and Methods

### Protein expression, purification and refolding

All proteins were expressed in a pET2B-T vector with an N-terminal His6 tag, followed by a TEV cleavage site, and the protein of interest. The amino acid sequences of the final purified protein constructs are given in **Table S1**. The expression and purification of all HP1α constructs were performed following previously described protocols.^23^ The Sgo1, CAF-1 and LBR constructs were all expressed in Rosetta BL21 (DE3) cells (New England Biolabs) at 37 °C for 4 hr where expression was induced with 0.5 mM isopropyl-β-D-thiogalactopyranoside (IPTG). The cells were lysed by sonication in 20 mM HEPES, pH 7.2, 300 mM KCl, 10% glycerol, and one Pierce protease inhibitor tablet (ThermoFisher Scientific). The lysed solutions were pelleted by centrifugation and the inclusion bodies were collected and washed twice with 20 mM HEPES, pH 7.2, 300 mM KCl, and 1% Triton X-100, followed by a final wash without Triton X-100. After the washes, 1 mL of dimethyl sulfoxide (DMSO) was added to the pellets and the samples were incubated on ice for 10 min. The proteins were extracted from the inclusion bodies using 20 mM HEPES, pH 7.2, 300 mM KCl, and 6 M guanidinium chloride and the mixture was nutated overnight at 4 °C, followed by centrifugation to collect the supernatant. The supernatant was then incubated with HisPur™ Ni-NTA resin (ThermoFisher Scientific), along with ∼6-8 mg of DNase I (New England Biolabs) and 10 mM MgCl_2_ at 4 °C for at least 1 hr. The resin was washed with a buffer containing 20 mM HEPES, pH 7.2, 300 mM KCl, 10% glycerol, 40 mM imidazole, and the proteins were eluted using 20 mM HEPES, pH 7.2, 300 mM KCl, 400 mM imidazole. The samples were then mixed with Tobacco etch virus (TEV) protease (prepared in house) at ∼ 1:20 TEV: protein mole ratio. Tris(2-carboxyethyl)phosphine (TCEP) was added at 1 mM final concentration and the samples were dialyzed overnight at 4 °C against buffer containing 20 mM HEPES, pH 7.2, 300 mM KCl, 0.5 mM TCEP. The dialyzed samples were collected and solid guanidinium chloride was added to the solution to a final concentration of 6 M. The pH was adjusted to ∼ 2 – 4 with trifluoroacetic acid (TFA) and the proteins were purified using reverse-phase high-performance liquid chromatography (HPLC) on an XBridge Peptide BEH C18 column (Waters) using a gradient solvent of acetonitrile with 0.1% TFA (30-40% for HP1α, 20-70% for LBR, 20-60% for Sgo1, 10-60% for CAF-1). The pure fractions were combined and the proteins were lyophilized for further storage. Final purity was assessed by analytical HPLC and ESI-TOF-MS. Note that for our LBR construct, which starts with a proline residue, the TEV cleavage was not performed, and the final construct retained its N-terminal hexahistidine tag.

The refolding procedure was similar for all proteins. Each lyophilized protein was dissolved in a buffer containing 20 mM HEPES, pH 7.2, 300 mM KCl, 6 M guanidinium chloride, 1 mM TCEP at a final protein concentration of 1 mg/mL. The solution was transferred into a 3,500 or 10,000 molecular weight cutoff SnakeSkin® dialysis tubing (depending on the protein), and dialyzed against a buffer containing 20 mM HEPES, pH 7.2, 1 M guanidinium chloride, 300 mM KCl, 1 mM TCEP at 4 °C for 6-8 hr, followed by overnight dialysis into 20 mM HEPES, pH 7.2, 300 mM KCl, 1 mM TCEP at 4 °C, followed by another buffer exchange in the same buffer for at least 4 hr. After refolding, the solution was concentrated to a volume of 200-500 μL, followed by the addition of 1 mL of fresh buffer. The sample was concentrated to a volume of 200-500 μL and the procedure was repeated two more times to ensure complete removal of any residual guanidinium.

### *In-vitro* phosphorylation and conjugation of Cy3 and Cy5 to various constructs

The *in vitro* phosphorylation of all HP1α constructs used casein kinase II (New England Biolabs) and was performed following our previously described protocol.^23^ For imaging experiments, each protein (HP1α constructs, Sgo1(400-527), Sgo1(456-527), or CAF-1(p150 subunit residues 182-295)) was labeled either with Cy3 or Cy5 using maleimide chemistry. Since the LBR constructs do not have any cysteine residues, we prepared LBR constructs with an additional cysteine at the C-terminus for imaging purposes. Each protein was incubated with 20x molar excess of Cy3 or Cy5 for 1 hr at ambient temperature with gentle rocking in a buffer containing 20 mM HEPES, pH 7.2, 300 mM KCl, 10 mM TCEP. Labeled HP1α was purified by reverse-phase HPLC, while all other labeled constructs were purified by size-exclusion chromatography. Successful conjugation was confirmed by LC-ESI-TOF-MS.

### Phase separation experiments

HP1α and pHP1α do not form condensates at high ionic strength, therefore the “no LLPS” controls were performed in 20 mM HEPES, pH 7.2, 300 mM KCl, and 1 mM TCEP buffer, while the phase separation experiments were performed in a buffer containing 20 mM HEPES, pH 7.2, 75 mM KCl, and 1 mM TCEP. Unless indicated otherwise, all phase separation samples contained 100 μM HP1α or pHP1α while the concentration of the protein binding partners was varied. After mixing, the samples were incubated on ice for at least 30 min and centrifuged at 200 x g for 10 min at 4 °C to separate the condensed and dilute phases. The total volume for each sample was 8 μL, allowing for 2 μL of the supernatant to be used for A280 measurements and another 2 μL to be used for analysis by SDS PAGE gel electrophoresis. The samples were analyzed on 15% acrylamide SDS PAGE gels, followed by Coomassie staining. Band intensity was quantified using ImageJ^43^ and normalized against the “no LLPS” control and concentration as determined by A280 measurements.

For microscopy experiments, larger volumes of sample were prepared (30 – 50 μL) and non-labeled proteins were mixed with 5-10% of labeled constructs. 7 – 8 μL of the sample was placed on a slide and dual-channel fluorescence microscopy experiments were performed using a Leica SP8 confocal microscope. The data was processed using Fiji ImageJ.^44^

### Binding assays

Native PAGE gel binding assays were performed using either HP1α CSD-CTE or HP1α CSD-CTE W174A with LBR (1-211), Sgo1 (400-527), Sgo1 (400-468), or Sgo1 (456-527). The samples were prepared in 20 mM HEPES, pH 7.2, 300 mM KCl, 1 mM TCEP, and 10% glycerol. The concentration of HP1α construct was 25 μM. The concentration of BP ranged from 0.01 μM to 125 μM. After mixing the HP1α construct with the BP, each sample was incubated at room temperature for 20 min and 10 μL of each sample was loaded onto a 5% TBE native gel. The gel running condition was 180 V for 30 min at 4 °C in 1X TBE running buffer. The gels were stained with Coomassie blue, imaged and ImageJ was used to determine the HP1α construct’s gel band intensities. To determine the fraction bound, the band intensity of each sample was divided by the intensity of a control band of HP1α alone run on the same gel. The data were analyzed using GraphPad Prism.

### NMR Experiments

The NMR samples were prepared in 20 mM HEPES, pH 7.2, 75 mM KCl, and 1 mM TCEP with 10% D_2_O. Samples contained ^15^N labeled 50 μM HP1α CSD-CTE with either 5 μM, 12.5 μM or 25 μM LBR (1-211), or 2.5 μM, 5 μM and 12.5 μM Sgo1 (400-527). An 800 MHz Avance Neo Bruker NMR spectrometer was used to collect the NMR data. The chemical shifts were referenced relative to the spectrometer frequency, with the water frequency at 4.717 ppm. Pulses were calibrated using the standard TopSpin protocol. HSQC experiments were performed with the pulse program fhsqcf3gpph,^45^ with 32 scans and a relaxation delay of 1.5 s. The following parameters were set for the ^1^H dimension: 13.583 ppm spectral width, frequency offset of 4.717 ppm, 2048 points and DQD acquisition mode. The following parameters were used for the ^15^N dimension: 36.00 ppm spectral width, frequency offset of 120 ppm, 128 points and States-TPPI acquisition mode. Data was processed in Bruker TopSpin and analyzed in POKY.^46^ The intensity cutoff for quantifying cross-peaks was 4 × *σ*_noise_ where *σ*_noise_ was the average noise level of the spectrum as calculated by POKY. The plotted intensity ratios represent *I_bound_* ⁄*I_free_* where *I_free_* represents the intensity of cross-peaks in the HP1α CSD-CTE spectrum, while *I_bound_* represents the intensity of the matching cross-peaks in the spectrum with binding partner. Error bars were calculated using the equation 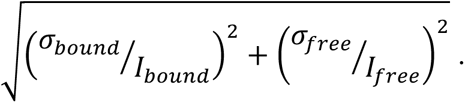 Data was plotted using GraphPad Prism. The data analysis took advantage of CSD-CTE homodimer assignments previously deposited under BMRB ID 53698.

### Coarse-grained molecular dynamics simulation protocol

Coarse-grained (CG) coexistence simulations were performed using the HOOMD-Blue 4.7 software package^47^ with slab geometry as described previously^48–50^. Proteins were modeled using the one-bead-per-residue HPS-Urry model^51^ with parameters for phosphorylated serine obtained from the HPS-PTM model^52^. For HP1α and pHP1α homodimers, the folded domains were treated as rigid bodies^53^ to prevent unfolding while allowing them to translate and rotate freely. Each CD was constrained as an independent rigid body, while two CSD monomers were constrained together as a single rigid body to maintain the dimerization interface. The disordered regions (NTE, hinge, and CTE) remained fully flexible, enabling interactions between all protein segments during the simulation. In simulations involving Sgo1 or LBR binding to the CSD dimer, the binding motifs PVVKI (Sgo1 residues 400-527) or VEVKL (LBR residues 1-211) were restrained to the CSD dimer interface. For HP1α (or pHP1α) simulations with Sgo1 or LBR at molar ratios of 1:1 or 1:2, the total protein concentration was maintained at ∼110 mg/mL, consistent with previous work^18,23^. Coexistence slab simulations were conducted for 5 μs in the canonical (NVT) ensemble at 320 K using a Langevin thermostat with friction coefficient γ = m_AA_/τ, where m_AA_ is the mass of each amino acid bead and τ is the damping time constant set to 1000 ps. The elevated temperature of 320 K was used based on our previous analysis^23^, which showed that this temperature ensures sufficient protein density in the dilute phase for accurate characterization of the coexistence behavior. The integration time step was 10 fs. The first 1 μs of each trajectory was discarded as equilibration before calculating density profiles and contact maps. Protein density was analyzed as a function of the z-coordinate to distinguish the dense and dilute phases. For contact analysis, two residues were considered in contact if their inter-bead distance was less than 1.5 times the arithmetic mean of their van der Waals radii. Simulation snapshots were visualized using VMD^54^.

## Results

### BPs can substantially modulate the condensation behavior of HP1α

Despite the growing literature focused on condensate formation and heterochromatin, HP1α on its own does not form condensates under physiological salt conditions *in vitro* even at high concentrations (**Fig. S2**).^10,13,23^ In our hands, N-terminal phosphorylation (pHP1α) can induce phase separation at concentrations of ∼ 75 μM in solutions containing 75 mM KCl (**Fig. S2**).^23^ The saturation concentration of HP1α required for condensate formation can be further reduced to the low μM range by the addition of nucleic acids, nucleosomes and nucleosome arrays.^7,16,35^ Here, we investigated the effect of BPs that specifically bind the CSD homodimer interface. For this purpose, we prepared recombinant CAF-1 (p150 subunit residues 182-295), LBR(1-211) and Sgo1(400-527), and tested a variety of BP and HP1α concentrations **(Fig. 2**). We labeled the BPs with the small fluorescent dye Cy5 and HP1α with Cy3, and monitored droplet formation and co-localization using fluorescent microscopy. Typically, the samples contained 5 – 10% of fluorescently labeled protein. On their own, none of the BPs underwent phase separation at the tested concentrations (**Fig. S3**). We chose to work at 75 mM KCl to compare with previous *in vitro* studies, the majority of which have been performed at this ionic strength.^10,23^

**Figure 2.**
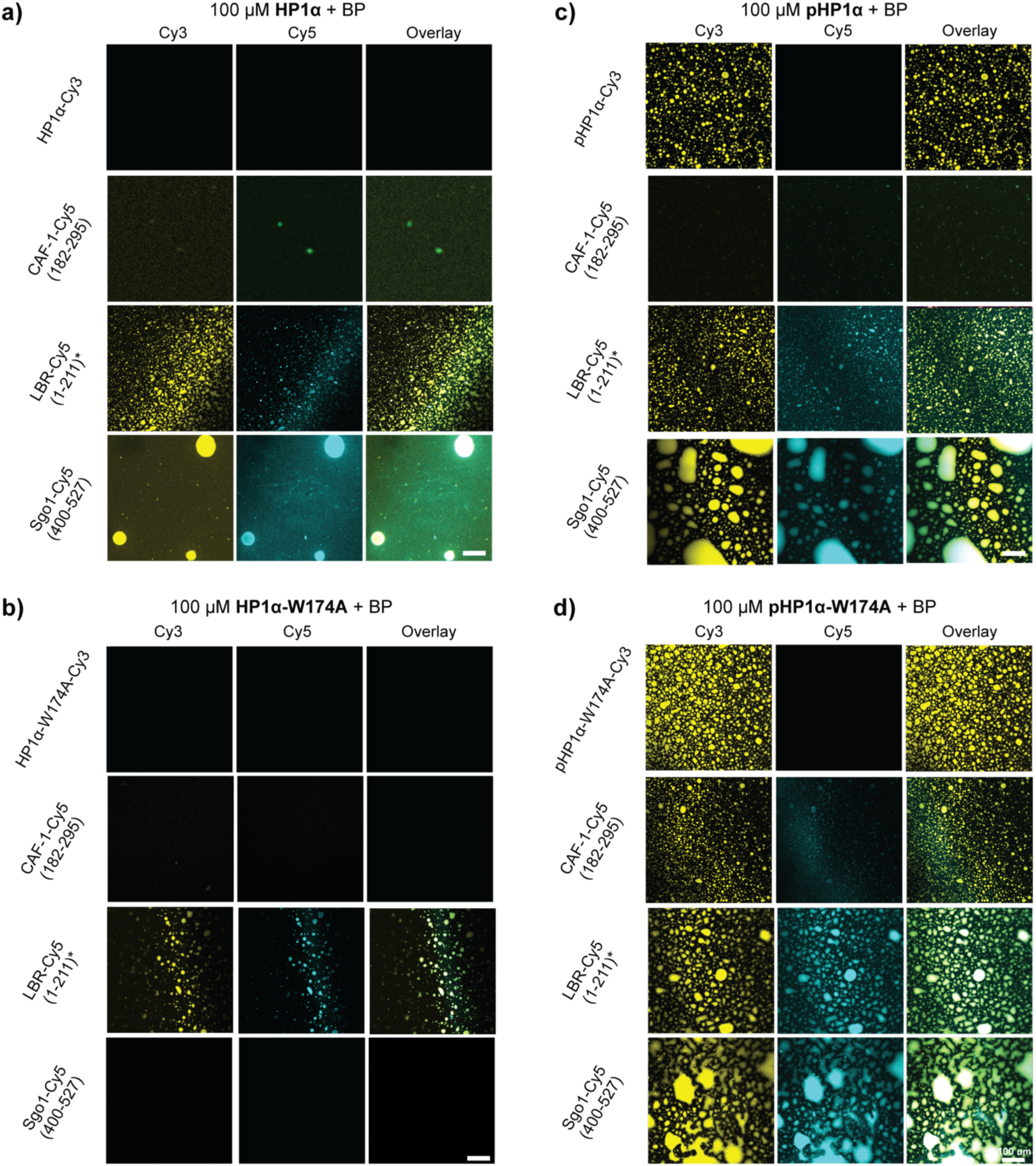
BPs modulate the condensation properties of HP1α. Fluorescence microscopy images of a) HP1α, b) HP1α with a W174A mutation, c) pHP1α and d) pHP1α with a W174A mutation in the presence of CAF-1(182-295), LBR(1-211) and Sgo1(400-527). Experiments were performed in 20 mM HEPES, pH 7.2, 75 mM KCl, and 1 mM TCEP. Samples contained 5% Cy3-labeled HP1α or pHP1α and were performed at a concentration of 100 μM protein to ensure robust phase separation of the pHP1α control. The concentration of BPs was 40 μM, including 5% Cy5-labeled protein. The scale bars represent 100 μm.

While CAF-1 had a small effect on HP1α phase separation, the effects of LBR and Sgo1 were much more dramatic (**Fig. 2a**). As assessed by microscopy, LBR and Sgo1 induced phase separation of HP1α at concentrations of HP1α as low as 5 μM (**Fig. S4, Fig. S5**). Thus, BPs can significantly enhance the condensation properties of HP1α and promote droplet formation at near physiological protein concentrations. We next wondered whether this effect depended on the specific HAC-CSD interactions. To answer this question, we prepared HP1α with a W174A mutation, which eliminated a crucial residue of the CSD homodimer binding interface, and assessed the ability of the BPs to modulate condensate formation (**Fig. 2b**). Interestingly, CAF-1 and Sgo1 no longer induced condensate formation even at 100 μM of HP1α, while LBR still retained this ability. This suggests that different BPs modulate phase separation through distinct mechanisms.

We next tested the effects of the BPs on the phase separation properties of pHP1α (**Fig. 2c**). At 100 μM pHP1α, the negatively charged CAF-1 segment significantly reduced condensate formation, while the positively charged LBR and Sgo1 preserved or enhanced this property. When pHP1α with the W174A mutation was used, robust condensation was still observed in all BP samples (**Fig. 2d**). Based on fluorescence alone, LBR and Sgo1 partitioned in pHP1α condensates, while CAF-1 appeared to be less efficient at entering the pHP1α condensates. These observations imply that electrostatic interactions are much more important in driving the phase separation properties of pHP1α compared to non-phosphorylated HP1α.

To provide a more quantitative view of the changes in condensation behavior as a function of BP, we prepared samples at different BP:HP1α ratios, tested for formation of condensates by microscopy, spun down the samples, collected the supernatant and analyzed the relative amount of HP1α in the dilute fraction by SDS-PAGE (**Fig. 3**). These experiments confirmed the qualitative observations by microscopy and provided additional details. In HP1α samples, for example, the concentration of HP1α in the supernatant did not change significantly as CAF-1 was added to the sample (**Fig. 3a**), suggesting that this BP does not have the ability to promote phase separation. In the presence of Sgo1 and LBR, the supernatant HP1α concentration decreased significantly, supporting the formation of condensates (**Fig. 3a**). When HP1α with the W174A mutation was used (**Fig. 3b**), only LBR appeared to promote condensate formation, similar to the observations by fluorescence microscopy. In the case of pHP1α, LBR and Sgo1 enhanced condensate formation irrespective of whether the HAC-CSD interactions were possible (**Fig. 3c,d**). CAF-1, on the other hand, disrupted the condensates when it could bind to pHP1α through the HAC motif, and did not affect the condensates when the HAC-CSD interaction was removed.

**Figure 3.**
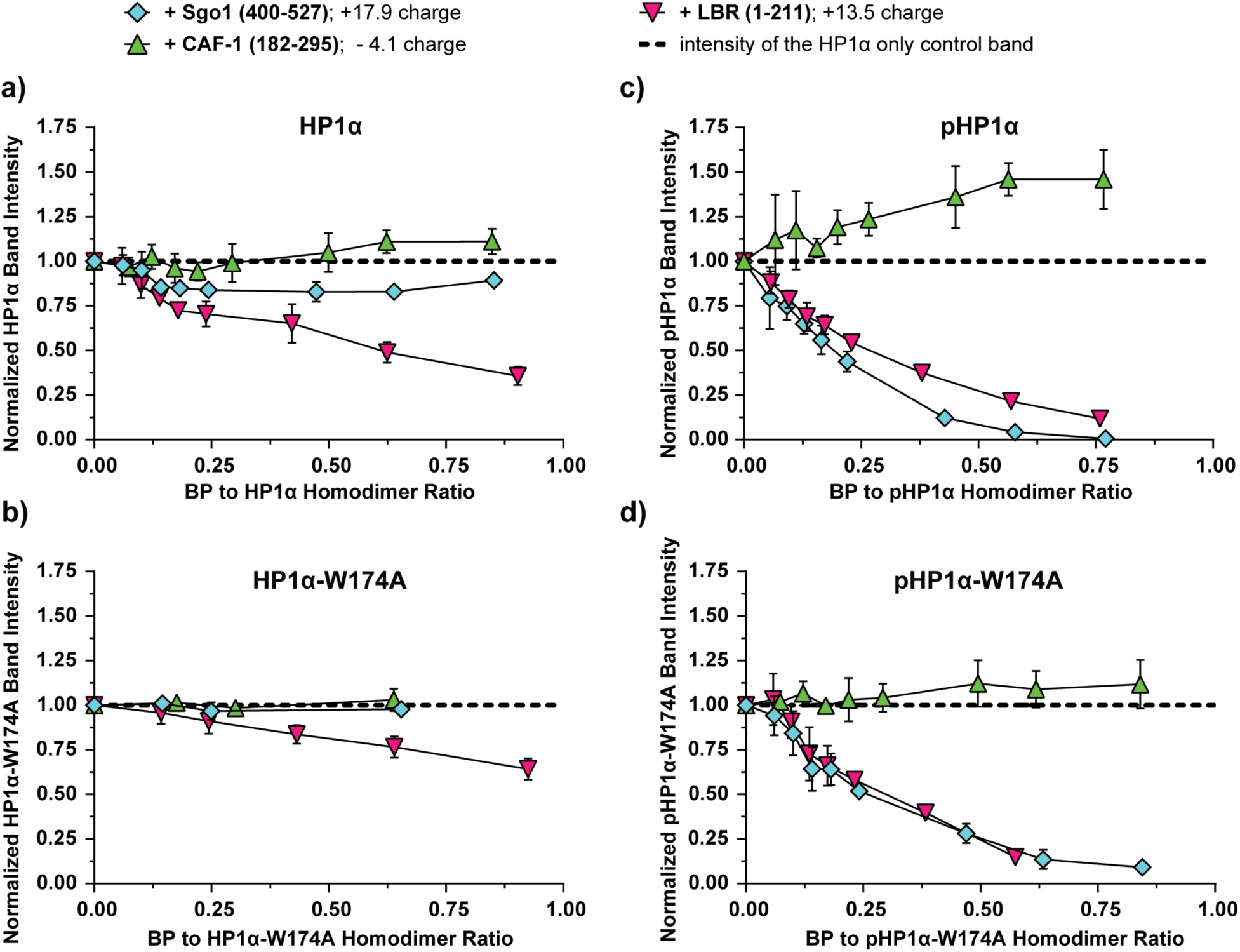
Quantitative analysis of the condensation propensity of HP1α as a function of BP ratio. Samples containing: **a)** HP1α, **b)** HP1α with a W174A mutation, **c)** pHP1α, and **d)** pHP1α with a W174A mutation, were mixed with varying ratios of BPs. The samples were centrifuged gently to separate the condensed and dilute phases and the relative amount of HP1α present in the dilute phase was analyzed by SDS-PAGE. The dashed line denotes the intensity of the HP1α only control band in each respective sample, blue diamonds represent samples containing Sgo1, green triangles denote samples containing CAF-1, and pink triangles indicate the presence of LBR. Experiments were performed in triplicate in a buffer containing 20 mM HEPES, pH 7.2, 75 mM KCl, and 1 mM TCEP, with 100 μM HP1α and varying concentrations of the relevant BP.

Taken together, our observations suggest that BPs can have distinct effects on the condensation properties of HP1α and pHP1α. In general, positively charged BPs can significantly enhance the phase separation of both HP1α forms, while negatively charged BPs display more nuanced behavior where CAF-1 has no effect on HP1α (i.e. does not induce phase separation) but disrupts condensate formation for pHP1α. The positively charged BPs, LBR and Sgo1, also appear to utilize HAC-CSD and electrostatic contacts to different extents in their interactions with HP1α. For example, LBR, which is less positively charged compared to Sgo1, can still promote HP1α phase separation even in the absence of HAC binding, while Sgo1 does not have this ability.

### BPs have different affinities to the HP1α CSD-CTE homodimer

Intrigued by the observed differences in condensation behavior exhibited by LBR and Sgo1 in the presence of HP1α and HP1α W174A, we wondered if those constructs may use distinct interaction modes to promote condensation. To answer this question, we first focused on the specific interactions between the HAC motifs present on LBR and Sgo1 and a CSD homodimer. LBR contains one HAC motif centered around the VEVKL sequence (residues 113 – 117), while the Sgo1 construct has two HAC motifs organized around residues PVVKI (451 – 455) and PAVAL (469 – 473).^32,33^ Binding studies between the CSD homodimer and various peptides and protein binding partners have typically been performed using Trp fluorescence, which does not require any additional labeling of the constructs.^55,56^ However, the CSD domain has two Trp residues, which may convolute the binding data analysis when the binding partners are long and have the possibility to interact at multiple sites. In addition, we wanted to test the affinity of the W174A CSD mutant to decouple specific HAC binding from other interactions. Therefore, we obtained relative binding affinity information using a native gel-based assay where 25 μM of the CSD homodimer were incubated with a range of BP concentrations and the amount of free CSD dimer was obtained from the relevant band intensity (**Fig. 4, Fig. S6**). While this experiment does not necessarily provide a specific K_d_, it allows us to assess the differences in relative affinities between the constructs. We also note that our CSD constructs contained the full CTE, i.e. we used CSD-CTE homodimers.

**Figure 4.**
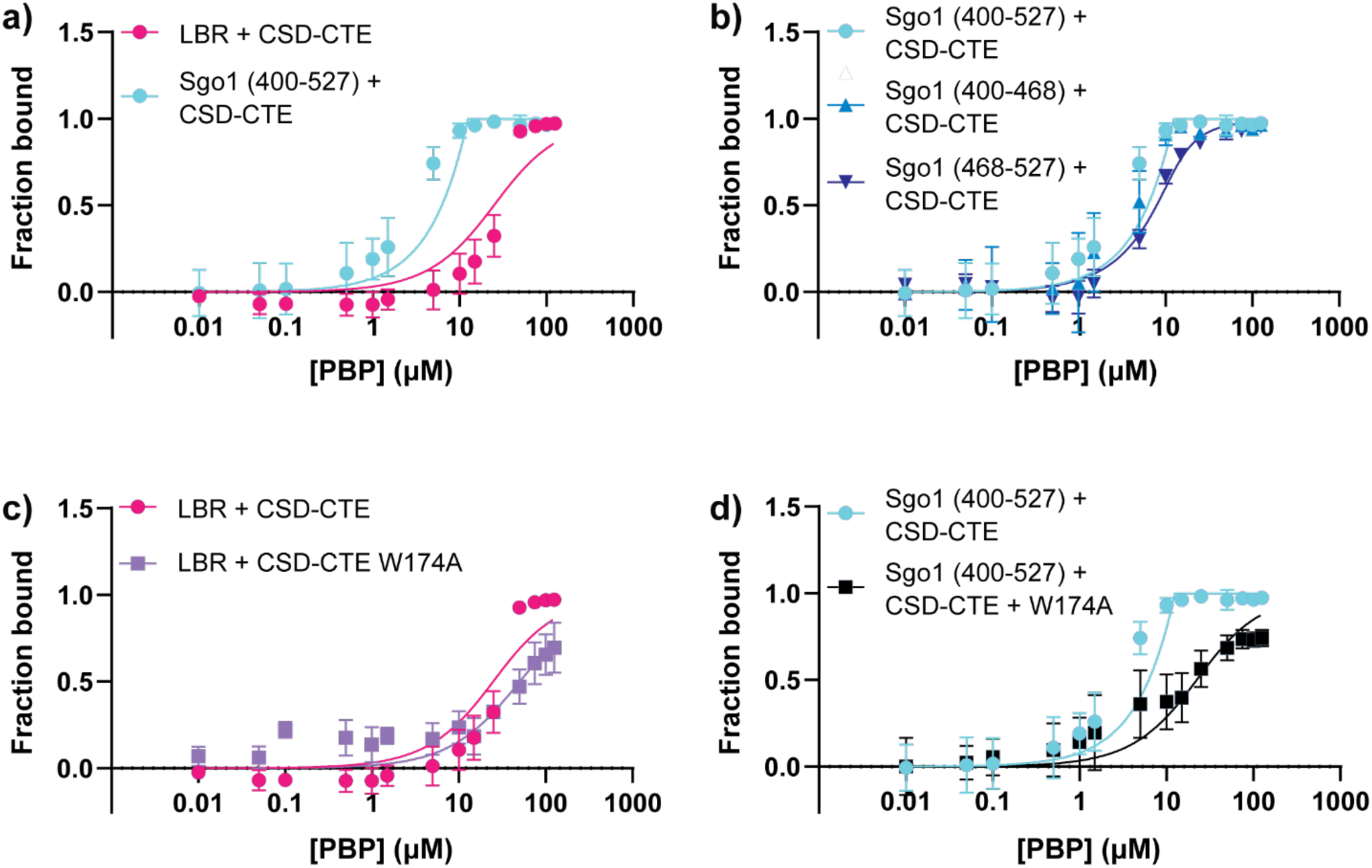
Relative binding affinity of various BP constructs. **a)** Comparison of the binding affinities of LBR(1 - 211) and Sgo1(400-527) to the CSD-CTE dimer. **b)** Comparison of the binding affinities of three different Sgo1 constructs: Sgo1(400-527) that comprises both HAC motifs; Sgo1(400-468) that contains the PVVKI HAC; and Sgo1(468-527) that contains the PAVAL HAC. **c)** Comparison of the binding affinity of LBR(1-211) to the CSD-CTE dimer or a dimer containing the W174A mutation. **d)** Comparison of the binding affinity of Sgo1(400-527) to the CSD-CTE dimer or a dimer containing the W174A mutation. All experiments contained 25 μM HP1α CSD-CTE (12.5 μM homodimers) and various concentrations of the protein binding partner (PBP). For representative native gels used to derive the binding affinity data, see **Fig. S6**.

Comparison of the data for Sgo1 and LBR indicates that Sgo1 binds the CSD-CTE homodimer with at least an order of magnitude lower K_d_ (**Fig. 4a**), while Sgo1 constructs containing only one of the two HACs behave very similarly to each other and to the construct containing both HAC motifs (**Fig. 4b**). These observations suggest that Sgo1 HACs have stronger binding affinity compared to the LBR HAC, and that they can bind interchangeably to the CSD-CTE homodimerization surface, at least within the limitations of the assay. Interestingly, the W174A mutation does not significantly impact the LBR interaction, while it reduces the binding affinity of Sgo1 by an order of magnitude. Altogether, these results suggest that Sgo1 interacts with the CSD-CTE homodimer with higher affinity and specificity, possibly enabled by the availability of two HACs and/or their more favorable sequence.

To further explore the interaction landscape between the two BPs and the CSD-CTE homodimer, we turned to solution NMR spectroscopy. We isotopically labeled the CSD-CTE homodimers with ^15^N and performed heteronuclear single quantum coherence (HSQC) titration experiments with increasing concentrations of unlabeled Sgo1(400-527) or LBR(1-211) (**Fig. 5**). The addition of increasing amounts of BPs lowered the intensity of the peaks in the HSQC experiments (**Fig. 5a,d, Fig. S7, S8**), and the effects were more pronounced for lower concentrations of Sgo1 compared to LBR. **Fig. 5** presents the analysis for 12.5 μM LBR and 5 μM Sgo1 where the overall intensity decreased but specific peaks with more significant intensity changes could be identified (with intensity change below one standard deviation from the average). In this context, both BPs induced significant intensity changes for HP1α residues at the CSD-CTE interface (E169/E170, L172 and H175) suggesting that they contact HP1α at the surface typically used for HAC interactions. In addition, they both contacted the neighboring loop at residues A129/T130 and C133. Finally, LBR induced significant intensity changes at W142, while Sgo1’s contacts extended a bit further to M137, F138 and L152. These patterns (**Fig. 5c,f**) suggest that both BPs contact HP1α through their HAC motifs and “wrap” around the neighboring surface of the CSD dimer, with Sgo1 making more extensive contacts at lower concentrations.

**Figure 5.**
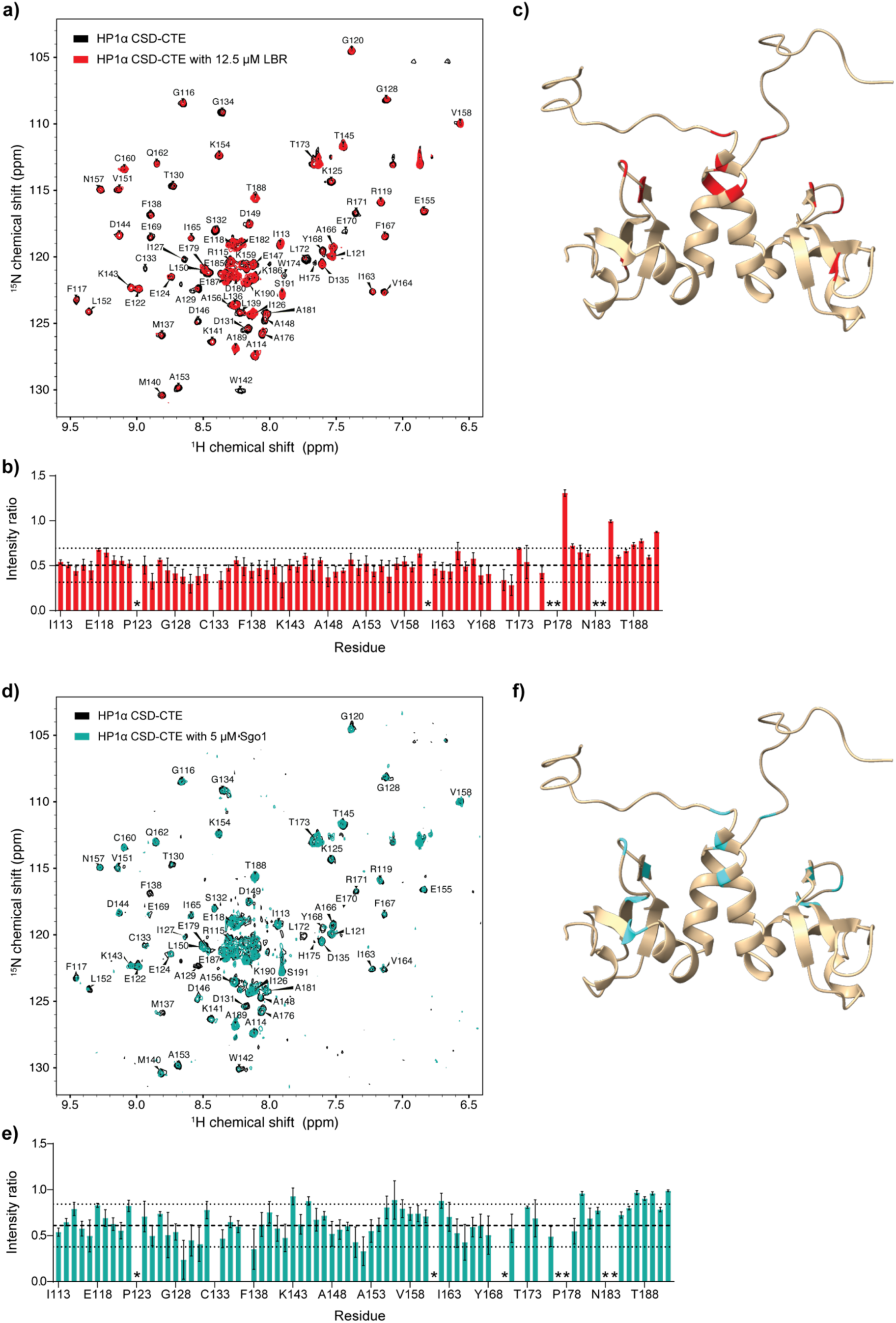
NMR studies of the CSD-CTE homodimer interactions with BPs. **a)** ^1^H-^15^N HSQC spectra of 50 μM CSD-CTE homodimer with and without 12.5 μM LBR (red and black, respectively). **b)** Analysis of the intensity changes of the CSD-CTE homodimer peaks in the presence of 12.5 μM LBR. The dashed line shows the average of the changes for all peaks, while the dotted lines show one standard deviation above and below. The error bars reflect the signal-to-noise in the spectra and peaks with signal-to-noise ratios of four or less were not included in the analysis. Residues with missing assignments or that did not meet the signal-to-noise criterion in the control spectrum are denoted with asterisks. **c)** Model of the CSD-CTE homodimer with residues showing significant intensity changes in the presence of LBR denoted in red. **d-f)** Analogous data for 5 μM Sgo1.

### BPs have different interaction maps with HP1α within condensates

Having explored the HAC interactions between Sgo1/LBR and the CSD-CTE homodimer, we next wondered how electrostatic contacts between the two BPs and the rest of the HP1α sequence may contribute to the condensation properties of the protein mixtures. To this end, we performed coarse-grained (CG) coexistence simulations based on the HPS-Urry model, which has been shown to successfully predict the relative saturation concentrations for condensate formation and to capture the residue-specific interactions within condensates.^18,23,51^ **Fig. 6** compares the relative dilute and dense phase concentrations for mixtures of HP1α and LBR(1-211) (**Fig. 6a**), and HP1α and Sgo1(400-527) (**Fig. 6b**). In these simulations, protein density was analyzed as a function of the z-coordinate to distinguish the dense and dilute phases, with the dilute phase concentration serving as a proxy for phase separation propensity, where lower concentrations indicate stronger driving forces for condensate formation. Consistent with our experimental observations, Sgo1(400-527) was less efficient at forming condensates with HP1α and its dilute phase concentration was higher (**Fig. 6c**) compared to the LBR construct. The interaction maps between BPs and HP1α provided further insights. LBR has a positively charged cluster between residues 60 and 100 (comprising the Arg/Ser (RS) repeat region^41^), which appears to be the main interaction hotspot on the protein, enabling interactions with the NTE, CD, CSD and the CTE of HP1α (**Fig. 6d**). On the other hand, the Sgo1 construct has a more promiscuous interaction map enabled by several interaction hotspots near the HAC motifs and the C-terminus. While both LBR and Sgo1 interact with the NTE, CD, CSD and the CTE of unmodified HP1α, the contact maps of pHP1α are dominated by contacts with the NTE (**Fig. S9**).

**Figure 6.**
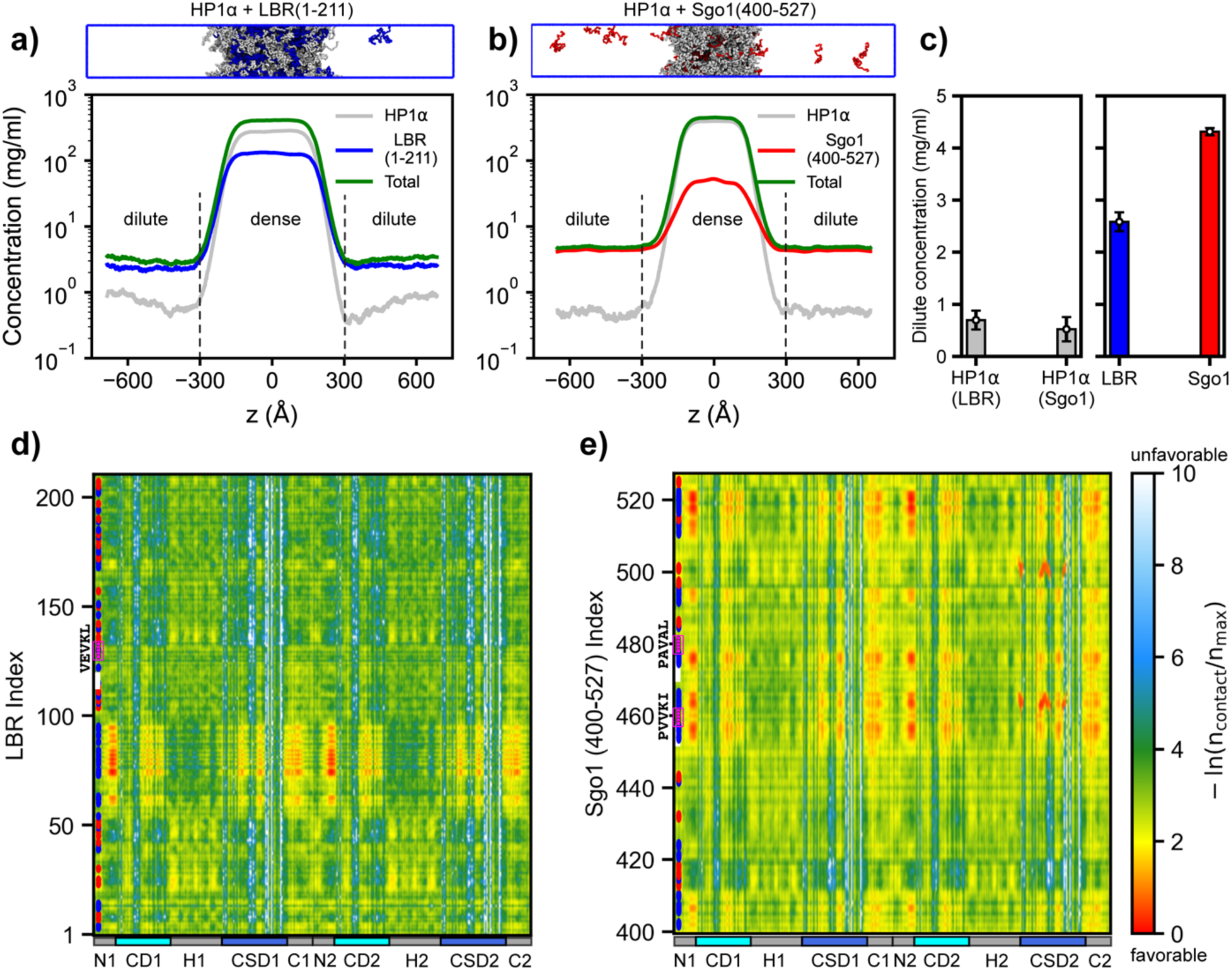
**Interaction maps of BP-HP1α condensates**. **(a, b)** Density profiles and representative condensate snapshots from coarse-grained (CG) coexistence simulations of HP1α with LBR (1–211) and Sgo1 (400–527) at a 1:1 mole ratio. The total protein concentration was maintained at ∼110 mg/mL, comparable to previous studies.^18,23^ Simulations were performed with the HPS-Urry model at 320 K. **(c)** Dilute-phase concentrations of HP1α and its binding partners in the CG simulations. Error bars represent block averages over five blocks. **(d, e)** Intermolecular contact maps between HP1α and LBR/Sgo1 within the condensates along the sequence of each protein. Blue and red dots on the y-axis denote the positions of positively and negatively charged residues, respectively. Binding motifs in LBR and Sgo1 are highlighted by magenta circles.

To explore the scenario where some BPs are bound to HP1α through the HAC motifs while others are not, we repeated the simulations but constrained 50% of the BP monomers to HP1α or pHP1α homodimers and assessed the effects on the dilute protein concentration and the interaction maps within the condensates (**Fig. S10**). Sgo1 was constrained through its PVVKI motif while LBR was constrained through its VEVKL motif. In this case, free monomeric Sgo1 was again less efficient at entering the condensates as evident by its higher dilute phase concentration. Interestingly, in this scenario, the interaction maps appear similar for each protein whether HP1α is phosphorylated or not (compare **Fig. S10f** and **h**, and **g** and **i**). This is particularly striking for Sgo1, where the predominant interaction with unphosphorylated HP1α is through the NTE (**Fig. S10g**), in stark contrast with the unbound scenario (**Fig. 6e**) where Sgo1 had an extensive interaction map with other HP1α domains. This suggests that bound BPs can screen some of the interaction sites on HP1α so that the NTE becomes the primary interaction site on HP1α. Our data suggests that bound Sgo1 is more efficient at doing so compared to bound LBR.

## Discussion

Despite its relatively small size of just 191 amino acids, HP1α displays a remarkably complex interaction network that underlies its ability to perform various functions in heterochromatin compartments, including compacting nucleosomes and DNA, orchestrating the installation of H3K9me2/3, and recruiting factors that are necessary for gene silencing and chromatin regulation.^1,17,24^ This versatility stems from its modular domain organization, which enables binding to H3 K9me2/3 modified histone tails through its CD domain,^4–6,16^ facilitates interactions with RNA and DNA through the positively charged hinge region,^7,16,25,26^ and allows the formation of homo- and heterodimers with other HP1 proteins through CSD-CSD interactions.^28^ In addition, the specificity and strength of these interactions can be modulated through post-translational modifications. For example, phosphorylation on the NTE, which is constitutively present in cells, favors specific H3 K9me2/3 binding over non-specific interactions with nucleic acids,^21,22,57^ while phosphorylation of serine residues in the hinge region dislocates HP1α from heterochromatin and targets it to centromeres.^17,32,58^ Furthermore, the formation of a new β-sheet interface upon CSD-CSD dimerization allows HP1α and HP1 proteins in general to interact with a multitude of other proteins, which facilitates the recruitment of silencing factors to heterochromatin, or, conversely, the recruitment of HP1α to specific locations in the nucleus.^31,34^ Our understanding of how these BPs regulate the interaction landscape of HP1α, and in turn, its properties and function, is largely incomplete. Here, we dissect the interactions between HP1α and three of its binding partners, CAF-1, Sgo1 and LBR, with a particular focus on their effect on HP1α phase separation. These proteins perform unique cellular functions and showcase the sequence diversity of HP1α BPs, thus allowing us to assess the common and distinct features that underlie their interaction networks with HP1α.

Comparison of the three BPs reveals that CAF-1 does not promote HP1α condensation and it diminishes the ability of pHP1α to form condensates. This is in contrast to the effects of LBR and Sgo1, which significantly increase the propensity of both HP1α and pHP1α to undergo phase separation. We attribute this effect to the overall negative charge of our CAF-1 construct (-4.1), which would clash with the overall negative charge of HP1α (-3.4) and pHP1α (-11.4). The elimination of HAC binding allows pHP1α to form condensates while CAF-1 does not enter these condensates very efficiently. This is similar to our previous observations with a shorter CAF-1 peptide (residues 210-238 which contains the HAC motif) suggesting that a significant portion of the CAF-1 sequence around the HAC motif disfavors HP1α/pHP1α condensation.^23^

CAF-1 has a well-known role in the deposition of histone proteins on newly synthesized DNA.^40^ In addition, it serves as a platform for the recruitment of key gene silencing factors, including HP1α, to aid the re-establishment of heterochromatin domains after DNA replication.^59^ In this context, it has been shown that the p150 (CHAF1A) subunit of CAF-1 can form nuclear bodies with liquid-liquid phase separation properties at the sites of integrated HIV long terminal repeats (LTRs).^60^ Interestingly, the deletion of residues in the HAC motif does not affect the ability of p150 to form nuclear bodies but leads to the dispersion of HP1α from the bodies. While our CAF-1 construct does not contain the residues important for p150-mediated nuclear body formation, our results are consistent with the view that CAF-1 recruitment of HP1α is primarily dominated by interactions involving the HAC motif rather than favorable electrostatic interactions with the rest of the HP1α sequence.

The HP1α interaction landscape of LBR appears to be much more complex. Our LBR construct can readily form condensates with HP1α and pHP1α and it can still do so in the absence of HAC interactions. This is reinforced by our binding studies, which show that the W174A mutation does not have a significant impact on the binding of LBR to the CSD-CTE homodimer. Our simulations suggest that the key interaction hotspot is the positively charged Arg/Ser (RS) repeat region between residues 60 and 100 on LBR, which can form contacts with the HP1α NTE and the overall negatively charged surfaces of the CD, CSD and the CTE. The interaction landscape refocuses upon HP1α NTE phosphorylation where contacts between the NTE and the RS domain on LBR dominate. This picture is different from our previous studies with a short LBR peptide (residues 105-124) which had a weak positive charge of +0.9 but could not induce the phase separation of pHP1α.^23^ Thus, the remainder of the LBR sequence, and in particular the RS region, appears to be important for the condensation properties of HP1α/pHP1α-LBR mixtures.

LBR functions at the inner nuclear membrane and it consists of a transmembrane domain that is involved in cholesterol synthesis and a nucleoplasmic N-terminal domain that helps tether heterochromatin to the nuclear lamina.^41^ Our construct (residues 1 – 211) comprises the soluble N-terminal domain which in turn consists of a TUDOR domain that binds histones H3 and H4,^61,62^ the positively charged RS repeats region that interacts with RNA and Lamin B,^61,63^ and the HAC sequence that is responsible for interactions with the CSD-CSD interface on HP1α.^30,33,34^ LBR forms microdomains at the nuclear lamina and it is currently unclear whether these domains form through a protein mediated condensation mechanism or involve lipid rearrangements at the inner nuclear membrane.^41,64^ Nevertheless, our work suggests that the N-terminal region of LBR can have a dramatic effect on the ability of HP1α and pHP1α to form condensates, suggesting a potential role of LBR in heterochromatin organization. While the HAC interaction does not appear to be the driving force for condensation in our two-component system, we speculate that it may play a more prominent role in a more complex scenario where HP1α has to compete for access to the RS LBR region with nucleic acids and other proteins such as Lamin B.

Finally, Sgo1 appears to rely both on electrostatic and HAC interactions to modulate the condensation properties of HP1α depending on the context. Our Sgo1 construct, which has a high net positive charge of +17.9, can efficiently induce condensates of both HP1α and pHP1α at physiological concentrations. For HP1α this property depends on the presence of the HAC interactions, while for pHP1α electrostatic interactions are dominant. This is similar to the behavior of a small Sgo1 peptide containing one of the HAC motifs (residues 446-466) and carrying a charge of +5.9.^23^ Nevertheless, our long Sgo1 construct appears to be less efficient at inducing HP1α phase separation compared to the longer LBR construct, despite its higher HAC affinity and higher net positive charge. This can be attributed to two factors. First, the higher affinity of the HAC interactions ensures that more Sgo1 is bound to the HP1α dimers which then positions the bound protein in such a way so that interactions with free Sgo1 are screened. This may weaken the multivalent interaction network necessary to build the two-component condensate. This is also evident from our NMR studies, which suggest that bound Sgo1 interacts more extensively with the CSD surface compared to LBR. In addition, Sgo1 has multiple but much smaller positively charged patches compared to LBR, which would create a more fragmented multivalent interaction network in the condensates.

Sgo1 is one of several HP1α BPs identified to have more than one HAC motif^34^ and in our binding studies, the two HAC regions appear to bind the CSD-CTE homodimer interface with similar affinities. This is noteworthy as a clear functional role for the Sgo1-HP1α interactions in mitosis has not been clearly established. HP1α also interacts with other centromeric proteins that contain HAC motifs, e.g. INCENP, so the presence of multiple HAC motifs on Sgo1 may allow it to compete more effectively for HP1α in the centromeric environment.^32^ In addition, while a specific role for Sgo1-HP1α condensation has not been established, previous literature suggests that the inner centromere has the properties of a biomolecular condensate that is scaffolded by the chromosome passenger complex (CPC, consisting of Aurora B kinase, INCENP, survivin and borealin).^65^ Interestingly, mitotically phosphorylated HP1α (which is phosphorylated at the hinge region) promotes CPC components condensation, while unphosphorylated HP1α increases the saturation concentration required for condensate formation.^65^ While our study focused on N-terminal phosphorylation, rather than hinge region phosphorylation, these observations are consistent with a view where the altered electrostatic properties of phosphorylated HP1α are important for its centromeric functions.

A final important point is that the HAC motifs themselves vary in strength and appear to be more complex than previously thought. Using Drosophila as a model system, Colmenares et *al.*, expanded the definition of the PXVXL/PXVXL-like HP1 binding motif to the so-called HP1 access code (HAC), which comprises a central 9-mer motif enriched in carefully positioned hydrophobic and positively charged residues surrounding a key Val or Ile residue.^34^ Additional positively charged or hydrophobic flanking resides expand the motif to 15 amino acids ensuring more robust binding to the CSD-CSD interface. Notably, strong HAC motifs are devoid of negatively charged residues. This framework explains why our LBR construct is a weaker binder than Sgo1 and why the W174A mutation in the CSD-CSD interface does not dramatically change its interactions with HP1α/pHP1α. Interestingly, our NMR data suggests that LBR still contacts the CSD-CSD dimer interface, but it forms less extensive interactions with the rest of the CSD surface. BPs with strong HAC motifs such as Sgo1, on the other hand, can use their extra hydrophobic and positively charged residues to more extensively “wrap” around the CSD-CSD dimer. This has important implications for the interaction network of HP1α as more extensive HACs may also lead to more screening of HP1α interaction surfaces, which in turn may modulate the competition between BPs for access to HP1α. In addition, our study illustrates the importance of studying longer (or ideally full-length) constructs of the BPs as short peptides centered around the PXVXL motifs may lack important HAC residues or may underestimate the role of electrostatic interactions. This may explain seemingly contradictory results where a short LBR peptide does not appear to promote pHP1α or pSwi6 (the fission yeast homolog) phase separation^10,23^ while our longer LBR construct can efficiently do so. Similarly, early peptide studies suggested that the Sgo1 PVVKI motif is a much stronger binder compared to the PAVAL motif but these studies did not explore the effects of the full 15-mer HAC motif sequences.^32^

## Conclusion

Overall, our studies demonstrate that HP1α BPs play an important role in modulating the condensation properties of HP1α and in regulating its complex interaction network. Each BP balances specific interactions through its HAC motif and electrostatic contacts to interact with HP1α and perform its unique functions in the cell in distinct nuclear locations. While the functional significance of some BP-HP1α interactions is still unknown, our study indicates that the diverse BP sequences have the ability to promote or enhance HP1α condensation, account for HP1α post-translational modifications, and screen or compete against other BPs for access to HP1α. This complexity suggests that HP1α, and possibly the other human paralogs HP1β and HP1γ, may have still unknown functions beyond those involved in gene and heterochromatin regulation. We look forward to future studies that will no doubt illuminate other important and interesting facets of these proteins.

## Data availability

All data is presented within the manuscript.

## Author contributions

C.H., R.B. and G.T.D. designed the research. C.H., R.B., T.D. and L.S.A. performed experiments, while T.M.P. performed the simulations. All authors analyzed the data and C.H., R.B., T.M.P., J.M. and G.T.D. wrote the manuscript with input from all authors.

## Acknowledgements

This work was supported by NIH grants R35GM138382 (G.T.D., C.H., R.B.), R35GM153388 (J.M.), K99GM159055 (T.M.P), and T32GM139795 (L.S.A), while T.D. was supported by a UCSD summer undergraduate research fellowship. We thank Dr. Xuemei Huang for assistance with NMR experiments. We are grateful for the computational resources provided by Texas A&M High Performance Research Computing (HPRC).

