## Supplementary information for "The role of electrostatic interactions in the phase separation of HP1α and its protein binding partners"

### Supplementary figures

**a)**

**Chromatin Assembly Factor-1 p150 subunit (CAF-1)**

**Charge at pH 7.2**

- 4.1 (Residues 182-295)

```
182       190       200       210       220       230       240
SKTEEEGVGCGGAGRRGDSQECSPRSCPELTSGPRMCPRKEQDSWSEAGGILFKGKVPMVVL

243    250       260       270       280       290
QDILAVRPPQIKSLPATPQGKNMTPESEVLESFPEEDSVLSHSSLSSPSSTSS
```

**b)**

Histidine Tag--**Lamin B Receptor (LBR)**

**Charge at pH 7.2**

+13.5 (Residues 1-211)

+4.4 (Residues 91-211)

```
    Histidine Tag
MKSSHHHHHHENLYFQ--

1        10        20        30        40        50        60
SPSRKFADGEVVRGRWPGSSLYYEVEILSHDSTSQLYTVKYKDGTELELKENDIKPLTSF

61       70        80        90        100       110       120
RQRKGGSTSSSPSRRRGSRSRSRSRSPGRPPKSARRSASASHQADIKEARREVEVKLTPL

121      130       140       150       160       170       180
ILKPFGNSISRYNGEPEHIERNDAPHKNTQEKFSLSQESSYIATQYSLRPRREEVKLKEI

181      190       200       210
DSKEEKYVAKFLAVRTFEVTPIRAKDLEFGG
```

**c)**

**Shugoshin-1 (Sgo1)**

**Charge at pH 7.2**

+17.9 (Residues 400-527)

+8.0 (Residues 400-468)

+14.8 (Residues 456-527)

```
400       410       420       430       440       450       460
SVTRPLAKRALKYTDEKETEGSKPTKTPTTTPPETQQSPHLSLKDITNVSLYPVVKIRRLSL

461       470       480       490       500       510       520
SPKKNKASPAVALPKRRCTASVNYKEPTLASKLRRGDPFTDLCFLNSPIFKQKKDLRRSK

521    527
KRALEVS
```

**Figure S1. Sequence of the HP1α BP constructs used in this study.** Positively charged residues (Lys and Arg) are highlighted in blue, while negatively charged residues (Asp and Glu) are highlighted in red. The HAC motifs in each protein are underlined with the central PXVXL or PXVXL-like sequence highlighted in yellow.

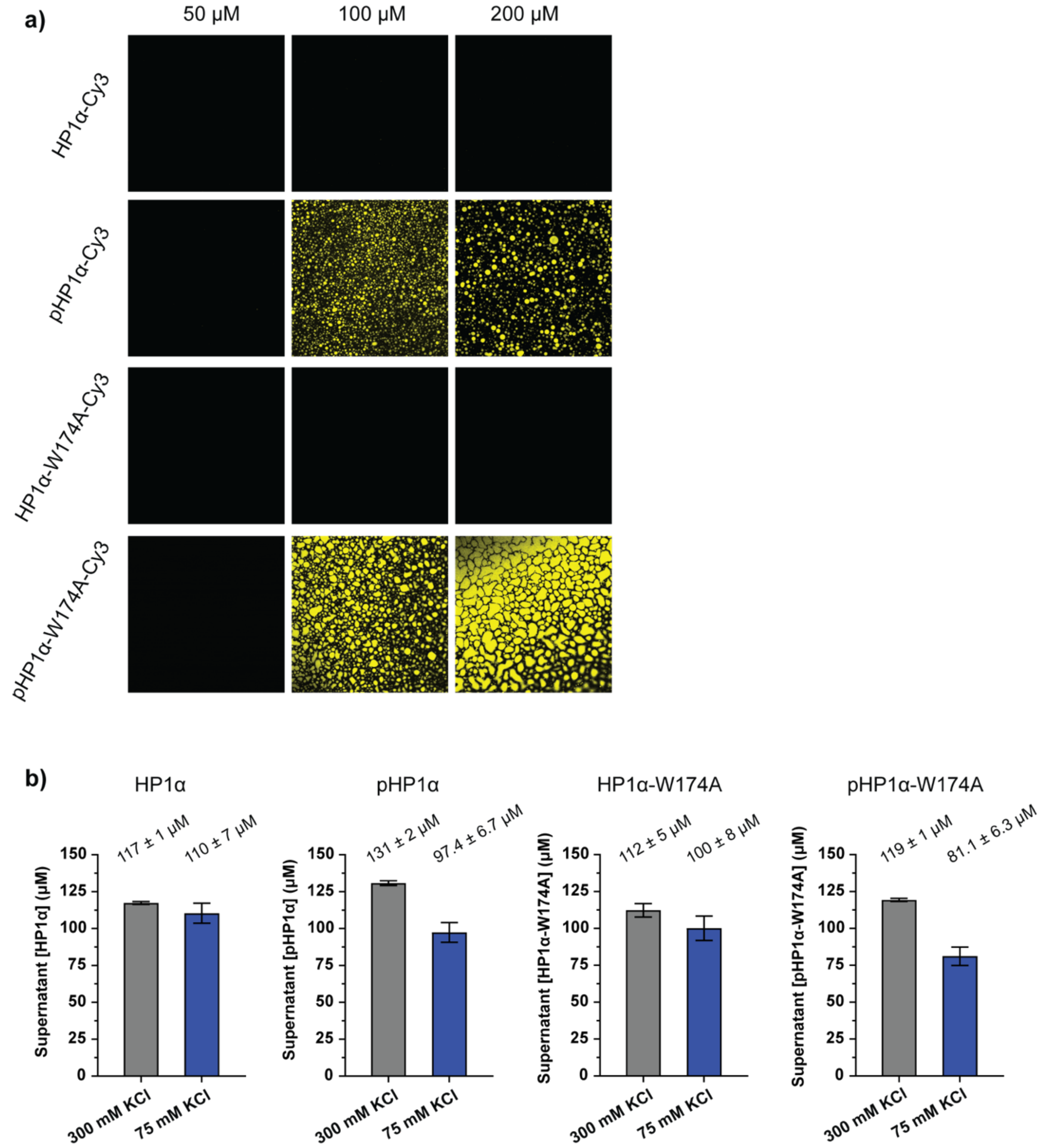


**Figure S2. a)** Phase separation behavior of relevant HP1α constructs at various concentrations. Experiments were performed in 20 mM HEPES, pH 7.2, 75 mM KCl, and 1 mM TCEP with 5% Cy3-labeled protein. **b)** Comparison of protein concentrations in the supernatant after samples were prepared at 300 mM KCl where phase separation does not occur and 75 mM KCl where phase separation is expected, followed by gentle centrifugation to separate the condense phase from the dilute phase. Concentrations were determined in triplicate based on A280 absorbance.

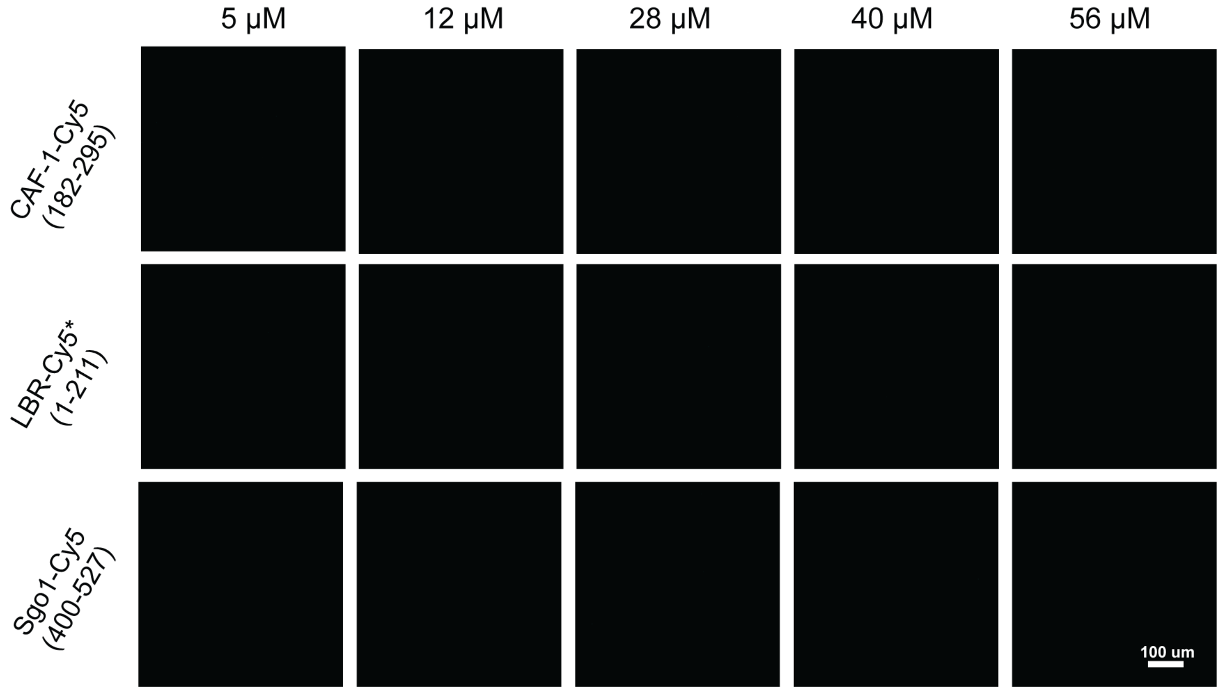


**Figure S3.** No condensates were observed in any of the BP samples at the tested concentrations. Experiments were performed in 20 mM HEPES, pH 7.2, 75 mM KCl, and 1 mM TCEP and samples contained 5% Cy5-labeled proteins.

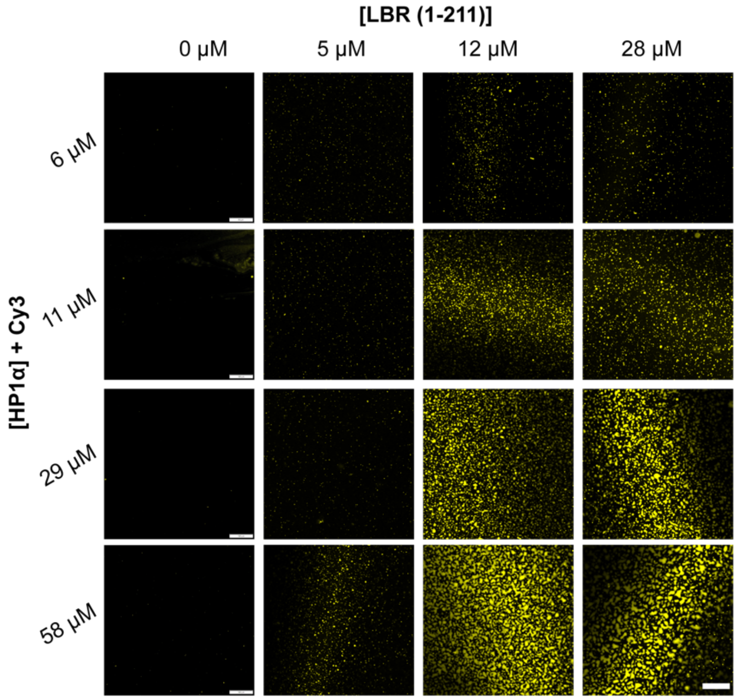


**Figure S4.** Fluorescence microscopy of samples containing HP1α-Cy3 and LBR(1-211) at various concentrations. Experiments were performed in 20 mM HEPES, pH 7.2, 75 mM KCl, and 1 mM TCEP and samples contained 5% Cy3-labeled protein. The scale bars denote 100 µm for the control samples (no LBR) and 50 µm for the rest of the samples.

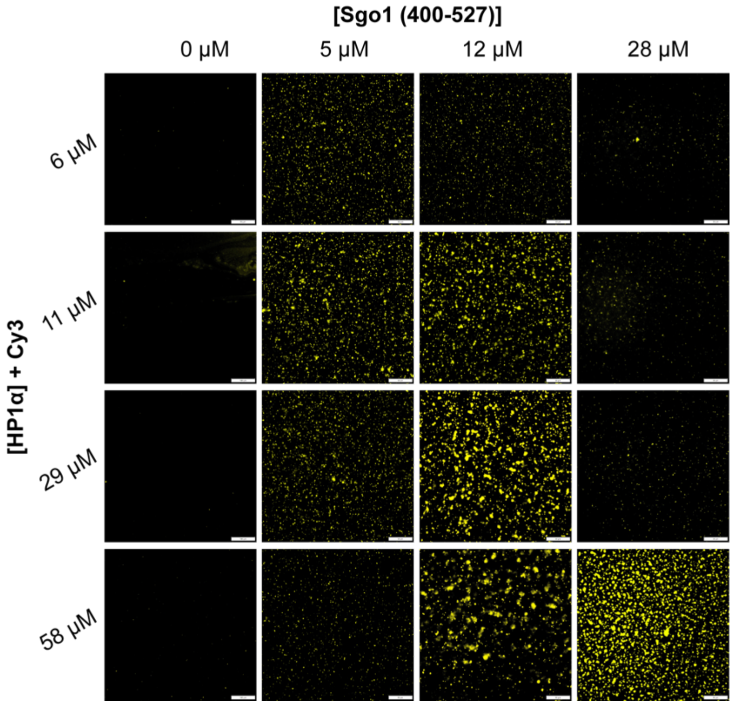


**Figure S5.** Fluorescence microscopy of samples containing HP1α-Cy3 and Sgo1(400-527) at various concentrations. Experiments were performed in 20 mM HEPES, pH 7.2, 75 mM KCl, and 1 mM TCEP and samples contained 5% Cy3-labeled protein. The scale bars denote 100 μm for the control samples (no Sgo1) and 50 μm for the rest of the samples.

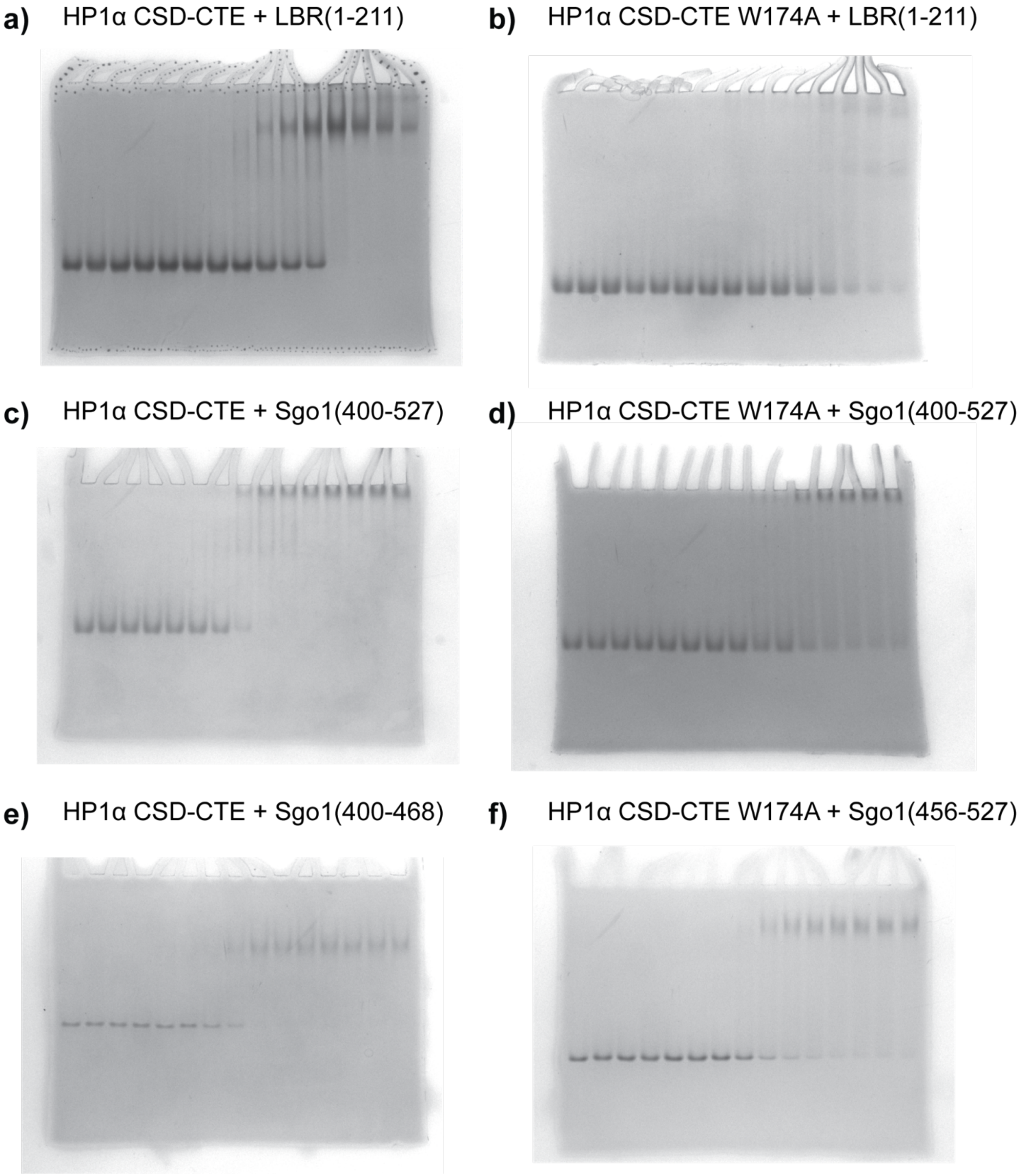


**Figure S6.** Representative native gels used to determine the relative binding affinities in Fig. 4. All experiments were performed with 25 μM HP1α CSD-CTE (12.5 μM homodimers) or HP1α CSD-CTE containing the W174A mutation and increasing concentrations of the BP as indicated in the figure.

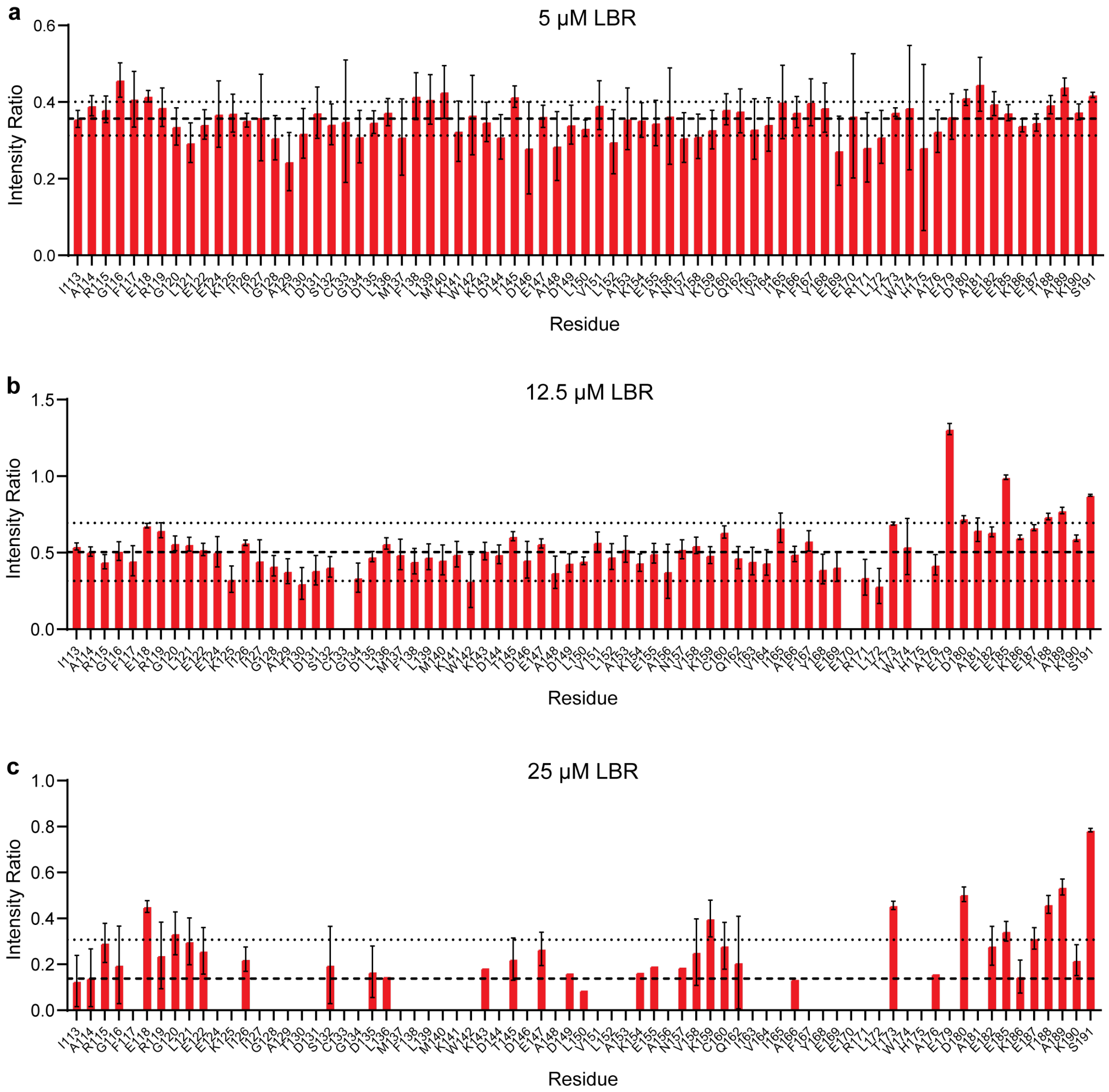


**Figure S7.** Intensity ratios of the CSD-CTE $^{1}H$-$^{15}N$ HSQC cross-peaks as a function of LBR(1-211) concentration.

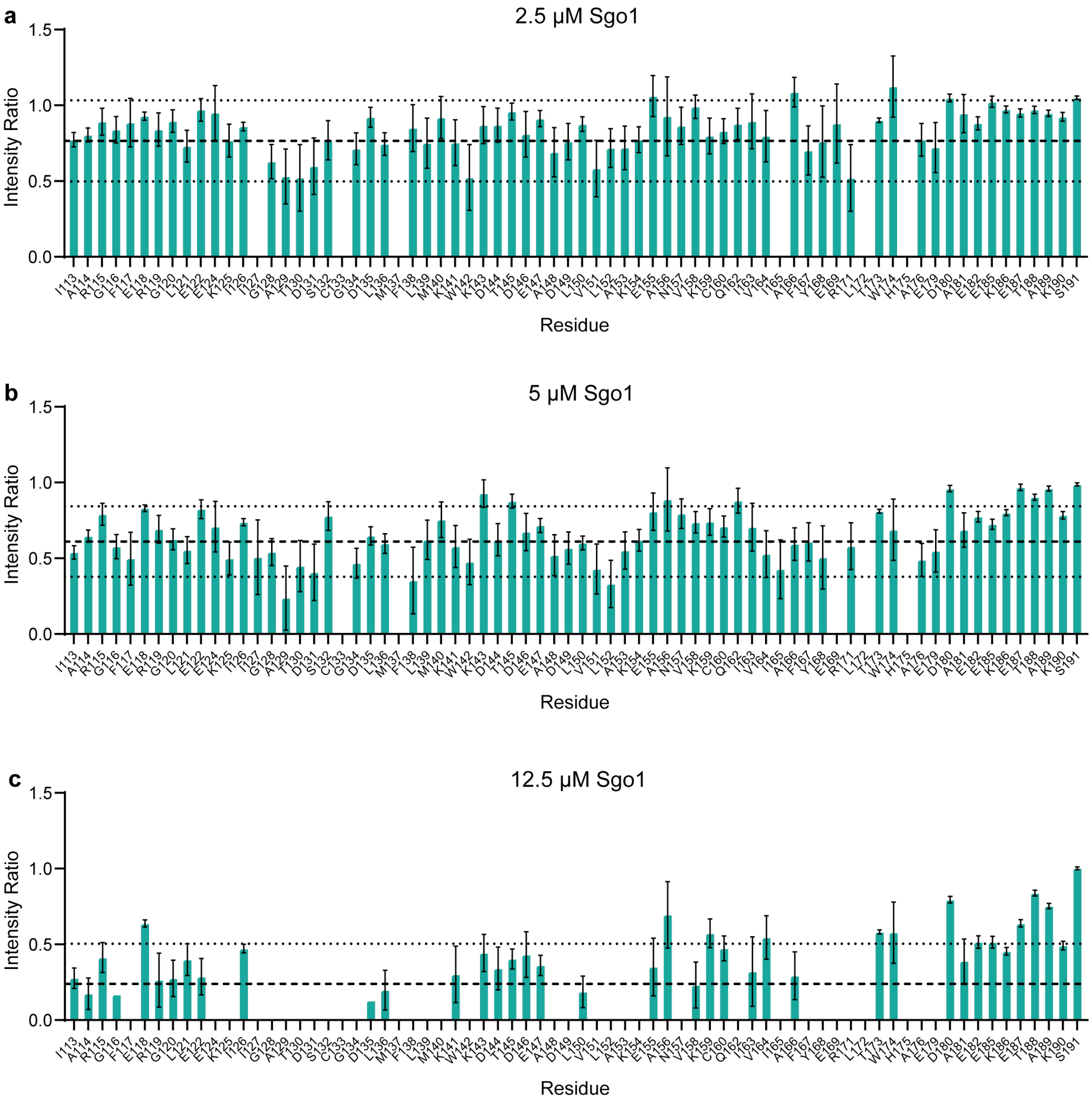


**Figure S8.** Intensity ratios of the CSD-CTE $^{1}H$-$^{15}N$ HSQC cross-peaks as a function of Sgo1(400-527) concentration.

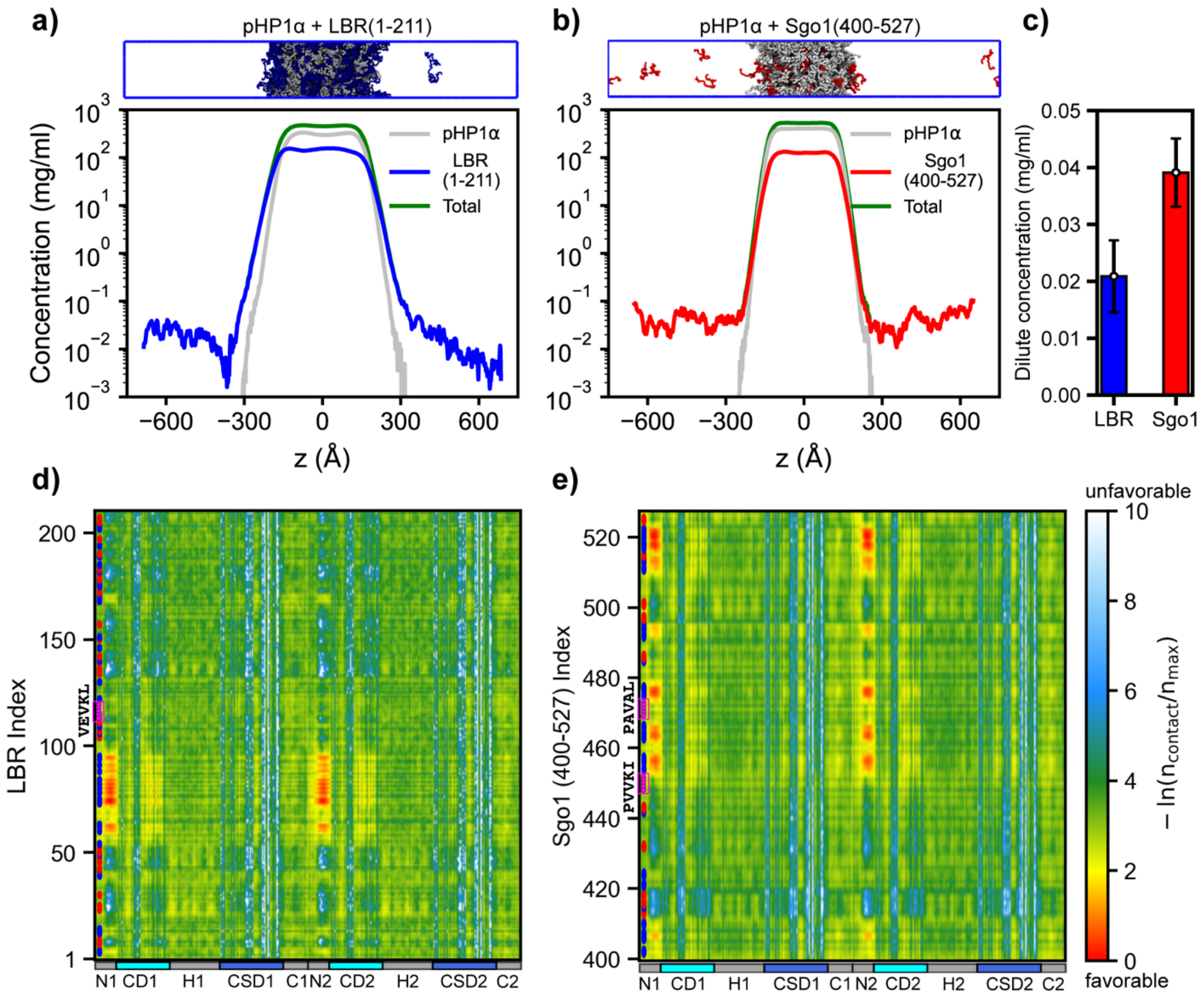


**Figure S9**. **Interaction maps of BP-pHP1α condensates.** (**a, b**) Density profiles and representative condensate snapshots from CG coexistence simulations of NTE-phosphorylated HP1α (pHP1α) with LBR (1–211) and Sgo1 (400–527) at a 1:1 mole ratio. The total protein concentration was maintained at ~110 mg/mL, consistent with previous studies (Her et al. 2022, Phan et at. 2024). Simulations were performed at 320 K using the HPS-Urry model, with phosphorylated serine parameters from the HPS-PTM model. (**c**) Dilute-phase concentrations of LBR and Sgo1 in the CG simulations. Error bars represent block averages over five blocks. (**d, e**) Intermolecular contact maps between pHP1α and LBR/Sgo1 within the condensates along the sequence of each protein. Blue and red dots on the y-axis denote the positions of positively and negatively charged residues, respectively. Binding motifs in LBR and Sgo1 are highlighted by magenta circles.

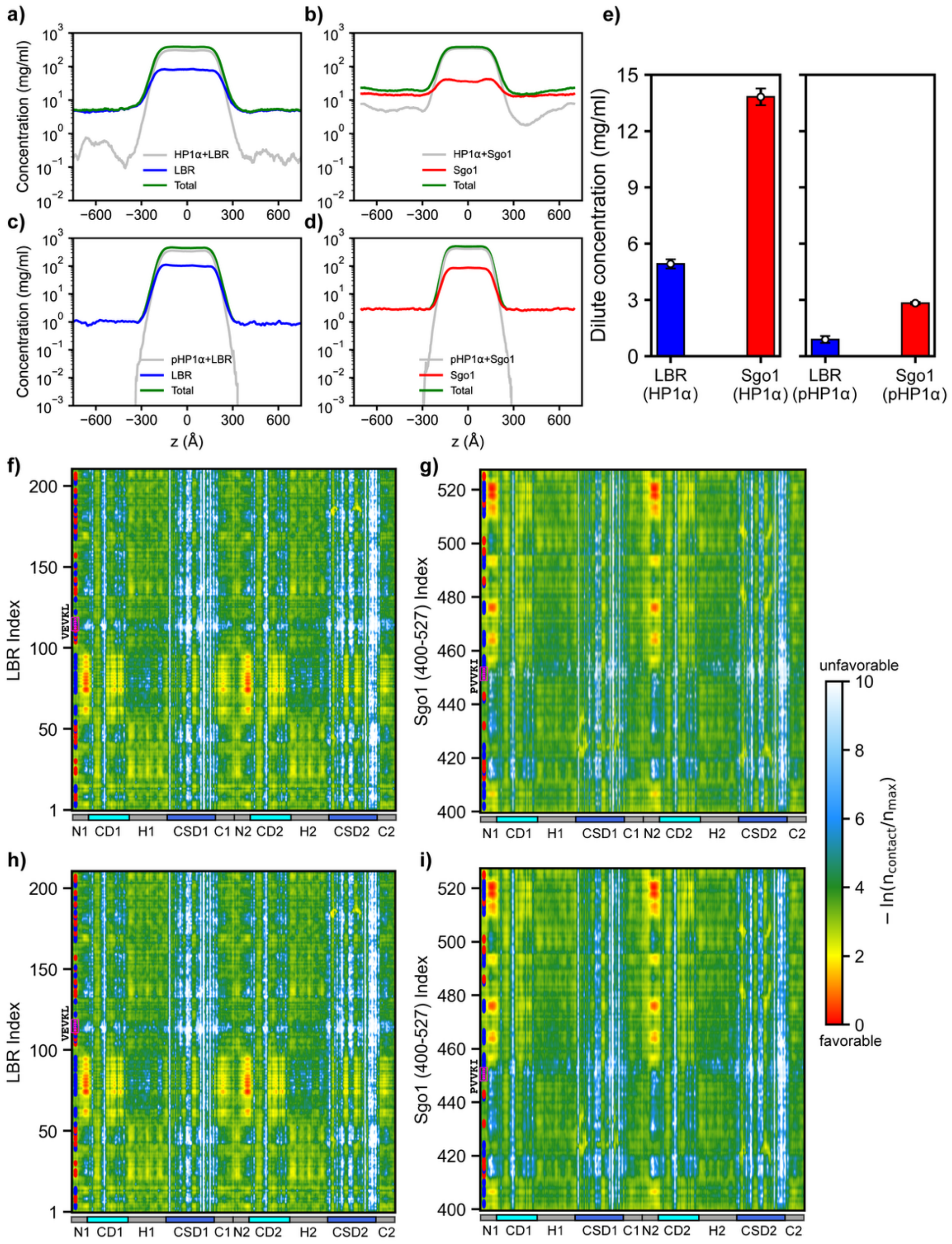


**Figure S10**. **Effect of BP restraints on the condensation behavior of HP1α and pHP1α.** (**a–d**) Density profiles from CG coexistence simulations at a 1:2 mole ratio of HP1α (**a, b**) or pHP1α (**c, d**) with binding partners. In these simulations, 50% of the BP is restrained to the HP1α CSD-CSD interface through the HAC motifs VEVKL (for LBR(1–211)) and PVVKI (for Sgo1(400–527)). (**e**) Dilute-phase concentrations of LBR and Sgo1 monomers when mixed with HP1α-LBR or

HP1α-Sgo1 complexes, or pHP1α-LBR or pHP1α-Sgo1 complexes. Error bars represent block averages over five blocks. (**f**) Intermolecular contact map between free LBR and HP1α within HP1α-LBR complexes in condensates. (**g**) Intermolecular contact map between free Sgo1 and HP1α within HP1α-Sgo1 complexes in condensates. (h) Intermolecular contact map between free LBR and pHP1α within pHP1α-LBR complexes in condensates. (i) Intermolecular contact map between free Sgo1 and HP1α within HP1α-LBR complexes in condensates. Blue and red dots on the y-axis denote the positions of positively and negatively charged residues, respectively. Binding motifs in LBR and Sgo1 are highlighted by magenta circles.

**Table S1. Various constructs used in this study**

| |
|---|
| HP1α, residues 1-191 (wild type):<br>**SGKKTKRTADSSSSEDEEEYVVEKVLDRRVVKGQVEYLLKWKGFSEEHNTWEPEKNLDCPELI**<br>**SEFMKKYKKMKEGENNKPREKSESNKRKSNFSNSADDIKSKKKREQSNDIARGFERGLEPEKI**<br>**IGATDSCGDLMFLMKWKDTDEADLVLAKEANVKCPQIVIAFYEERLTWHAYPEDAENKEKETA**<br>**KS** |
| HP1α-W174A, residues 1-191:<br>**SGKKTKRTADSSSSEDEEEYVVEKVLDRRVVKGQVEYLLKWKGFSEEHNTWEPEKNLDCPELI**<br>**SEFMKKYKKMKEGENNKPREKSESNKRKSNFSNSADDIKSKKKREQSNDIARGFERGLEPEKI**<br>**IGATDSCGDLMFLMKWKDTDEADLVLAKEANVKCPQIVIAFYEERLTAHAYPEDAENKEKETA**<br>**KS** |
| CAF-1, p150 subunit residues 182-295:<br>S**KTEEEGVGCGGAGRRGDSQECSPRSCPELTSGPRMCPRKEQDSWSEAGGILFKGKVPMVVLQ**<br>**DILAVRPPQIKSLPATPQGKNMTPESEVLESFPEEDSVLSHSSLSSPSSTSS** |
| Sgo1, residues 400-527:<br>S**VTRPLAKRALKYTDEKETEGSKPTKTPTTTPPETQQSPHLSLKDITNVSLYPVVKIRRLSLS**<br>**PKKNKASPAVALPKRRCTASVNYKEPTLASKLRRGDPFTDLCFLNSPIFKQKKDLRRSKKRAL**<br>**EVS** |
| Sgo1, residues 400-468:<br>S**VTRPLAKRALKYTDEKETEGSKPTKTPTTTPPETQQSPHLSLKDITNVSLYPVVKIRRLSLS**<br>**PKKNKAS** |
| Sgo1, residues 456-527:<br>S**RRLSLSPKKNKASPAVALPKRRCTASVNYKEPTLASKLRRGDPFTDLCFLNSPIFKQKKDLR**<br>**RSKKRALEVS** |
| LBR, residues 1-211 (with 6xHis-tag on N-terminus):<br>MKSSHHHHHHENLYFQS**PSRKFADGEVVRGRWPGSSLYYEVEILSHDSTSQLYTVKYKDGTEL**<br>**ELKENDIKPLTSFRQRKGGSTSSSPSRRRGSRSRSRSRSPGRPPKSARRSASASHQADIKEAR**<br>**REVEVKLTPLILKPFGNSISRYNGEPEHIERNDAPHKNTQEKFSLSQESSYIATQYSLRPRRE**<br>**EVKLKEIDSKEEKYVAKELAVRTFEVTPIRAKDLEFGG** |
